# A versatile reporter system to study cell-to-cell and cell-free bovine leukemia virus infection

**DOI:** 10.1101/2024.02.26.581980

**Authors:** Florencia Rammauro, Martín Fló, Federico Carrión, Claudia Ortega, Francesca Di Nunzio, Alexander Vallmitjana, Otto Pritsch, Natalia Olivero-Deibe

## Abstract

Bovine leukemia virus (BLV) is a B-lymphotropic oncogenic retrovirus of the genus *Deltaretrovirus* that infects dairy cattle worldwide and is the causative agent of enzootic bovine leukosis. BLV demonstrates remarkably low efficiency in infecting cells via free viral particles derived from infected B cells, as virions are rarely detected in the bloodstream of infected cattle. However, transmission efficacy significantly increases upon the establishment of direct cell-to-cell interactions. Syncytium formation assays are the main tool in the study of BLV infectivity. Although this traditional method is highly robust, the complexity of visually counting syncytia poses a significant technical challenge. Using lentiviral vectors, we generated a stable reporter cell line, in which the GFP reporter gene is under the control of the full-length BLV-LTR. We have demonstrated that the BLV-Tax protein and histone deacetylases inhibitors, such as VPA and TSA, can transactivate the BLV LTR. Upon co-culturing the reporter cell line with BLV-infected cells, fluorescent syncytia can be visualized. By implementing automated scanning and image acquisition using a confocal microscope, together with the development of an analysis software, we can detect and measure single GFP cells and fluorescent multinucleated cells. In summary, our reporter cell line, combined with the development of analysis software, is a useful tool for understanding the role of cell fusion and cell-free mechanism of transmission in BLV infection.

## INTRODUCTION

Bovine leukemia viru*s* (BLV) is an oncogenic deltaretrovirus that infects cattle worldwide and causes Enzootic bovine leukosis (EBL). BLV causes persistent subclinical infection (60% of the infected animals are asymptomatic, about 30% develop persistent lymphocytosis, and about 5-10% develop an aggressive tumor pathology called lymphosarcoma, which frequently causes the death of these animals [1], [2]. BLV infection leads to high economic losses in dairy and beef industries, related to mortality caused by lymphosarcoma [3]; negative impact on production parameters [4],[5] ; immunological alteration and secondary infections [6]; and restriction on the international trade of live cattle, semen, and infected embryos [7], [3], [8].

In contrast to retroviruses like HIV, deltaretroviruses such as BLV or human T cell leukemia virus (HTLV) exhibit remarkably low efficiency in infecting cells via free viral particles derived from B or T lymphocytes, respectively, as these virions are seldom detected in the bloodstream of infected hosts [9], [10]. Thus, efficient BLV transmission from a seropositive bovine to a susceptible counterpart primarily occurs via cell-containing fluids, such as blood, semen, milk, colostrum, saliva, or mucus, through horizontal or vertical routes [9].

Transmission efficacy substantially increases upon the establishment of direct cell-to-cell interactions. This mode of spread not only facilitates rapid viral dissemination but may also promote immune evasion and influence disease progression [11], [12], [13]. Cell-cell fusion (syncytium formation) has been identified as an alternative pathway for cell-to-cell transmission [13, 14].

Due to the intrinsic difficulty of infecting cells with cell-free BLV virions, syncytium formation assays are the main tool in the study of BLV infectivity [15–17]. This assay involves co-culturing an “acceptor” cell line, typically CC81 with a cell line persistently infected with BLV, such as FLKBLV, as the “donor”. The syncytia (multinucleated cells) form by fusion between an infected and an uninfected cell due to co-culture [16, 18]. After co-cultivation, the cells are commonly stained with Giemsa, and the number of syncytia is visually counted. Although this traditional method is highly robust, the complexity of visually counting syncytia poses a significant technical challenge, especially in high-throughput screening assays.

Reporter cell lines capable of detecting infectious viruses have been developed utilizing a BLV long terminal repeat (LTR) as a promoter and GFP or Luciferase as the reporter gene. Infected cells can be monitored using fluorescence microscopy or luminescence. These reporter cells offer greater sensitivity and quantification compared to traditional methods [19–22].

The BLV LTR promoter bears many regulatory sites and is responsible for virus integration and replication. It comprises by three distinct regions: the U3, R, and U5. The U3 region, contains several critical cis-acting elements besides to the CAAT box, TATA box, and transcription start site [23, 24]. The primary regulatory elements are three copies of an imperfectly conserved 21-bp sequence called the Tax-responsive element (TxRE). The TxREs are essential for the promoter’s responsiveness to the Tax transactivator protein. The BLV LTR has been suggested as a gene promoter under Tax activation in mammalian cells [19]. A glucocorticoid-responsive element (GRE) responds to dexamethasone in the presence of glucocorticoid receptors and Tax [25]. Additionally, there are multiple binding sites for several transcription factors, including two AP-4 sites, a glucocorticoid response element (GRE), and a PU.1/Spi-B–binding site. In the U5 region, a binding site for the interferon-responsive factor is present. These binding sites regulate BLV transcription, dependent on or independently of BLV-Tax expression [21].

Moreover, BLV transcriptional activity is influenced by the acetylation and methylation of these binding sites [26].

Using lentiviral vectors, we generated a stable reporter cell line to measure BLV infectivity, in which the EGFP reporter gene was expressed under the control of the full-length BLV-LTR. We have demonstrated that the Tax protein of BLV and histone deacetylases inhibitors (HDACi) such as, VPA and TSA, can transactivate the BLV LTR. When cultured with BLV-infected cells, the reporter cell line forms fluorescent syncytia. By integrating automated scanning and image acquisition using a confocal microscope, and developing an analysis software, we can detect single GFP cells and fluorescent multinucleated cells with two or more nuclei. Our results show that this reporter cell line is sensitive and, when combined with automated analysis, provides a useful tool to study both cell-to-cell and cell-free infection.

## MATERIAL AND METHODS

### Cell culture

HEK293T (ATCC CRL-3216) and CC81 (feline cell line transformed by mouse sarcoma virus ECACC 90031403) cells were grown in Dulbecco’s modified Eagle’s medium (DMEM)-high glucose (GlutaMAX, Gibco) supplemented with 1% penicillin/streptomycin and 10% fetal bovine serum (Gibco, USA). Persistently infected FLKBLV (Fetal lamb kidney, DSMZ-ACC 153) and BL3.1 (bovine B-lymphosarcoma, ATCC CRL-2306) cells were grown in Roswell Park Memorial Institute (RPMI) 1640 Medium-high glucose (GlutaMAX, Gibco) supplemented with 10% inactivated fetal bovine serum, 1% penicillin/streptomycin, 1% Sodium pyruvate. All cell lines were incubated at 37ºC in a 5%CO_2_ humidified atmosphere.

### Plasmids

DNA sequences encoding the full BLV Tax protein (GenBank, access number: EF600696) was synthesized and cloned in pcDNA3.1(+)-C-DYK by Genescript. The resulting plasmid, named pcDNA3.1BLVTAX-FLAG, encodes FLAG-tagged BLV Tax. pTrip lentiviral vector (pTripCMVGFP) [27], in which GFP are under the control of CMV promoter was used as a reporter or transfer vector. The CMV promoter was substituted with the 5’ LTR of BLV (531 pb, GenBank, access number: EF600696) to generate the pTripLTRBLVGFP vector. 5’ LTR of BLV was synthesized and cloned in (pTripCMVGFP) by Genescript.The commercial plasmids Lenti DR8.75 (Addgene #22036) and pMD2.G (Addgene #12259) were employed as packaging and envelope vectors, respectively.

### Transfection of HEK293T cells to obtain lentiviral particles (LVs)

HEK293T cells were co-transfected with the following plasmids: pTripLTRBLVGFP or pTripCMVGFP, DR8.74, and pMD2.G, to obtain lentiviral particles (LVs) with the VSV G protein on the surface and coding for GFP under the LTR of BLV (LVs LTRBLV-GFP) or under the CMV promoter as a control (LVs CMV-GFP). The day before transfection, 2.5x10^6^ HEK293T cells were seeded in 10 mL of complete medium (DMEM-GlutaMAX, 10% FBS and 1% PenStrep) into 100 mm petri dishes. PEI at 1 mg/mL (Polysciences) was used as the transfection agent. DNA:PEI complexes in a 1:10 ratio were prepared using 10 μg of pTripLTRBLBGFP or pTripCMVGFP, 6.5 μg of DR8.74, and 3.5 μg of pMD2.G, and added to the cells. After 12-18 h post-transfection, the medium was substituted with fresh medium. Forty-eight hours post-transfection, supernatant containing LVs was harvested and filtered by 0.45 µm and stored in aliquots at –80 ºC.

### Transduction for the establishment of the stable reporter cell line

The day before transduction, 0.5x10^5^ CC81, or FLKBLV cells per well were seeded in a 24-well plate to achieve a monolayer with 70% confluency. For transduction, the culture medium was removed, and 250 µL of LVs (LVs LTRBLVGFP or LVs CMVGFP) or cell culture medium (in negative control cells) together with 10 µL of polybrene (250 µg/mL) were added to each well. An additional 250 µL of cell culture medium was added to all wells, and the plate was incubated at 37°C and 5% CO_2_ for 60 h. GFP expression was evaluated by means of an epifluorescence microscope and images were inspected using ImageJ. A frozen stock of transduced cells was generated, and some cells were expanded for further experiments. The stable cell lines generated were designated CC81LTRBLVGFP or FLKLTRBLVGFP.

### Transient transfection of CC81LTRBLVGFP cells with plasmids encoding BLV Tax protein

1x10^5^ CC81LTRBLVGFP cells per well were seeded in a 12-well plate for 24 hours, and the cell culture medium was changed 3 hours before transfection. The PEI transfection reagent at 1 mg/mL was prepared in OptiMEM medium and added to a tube containing pcDNA3.1BLVTAX-FLAG or pcDNA3.1 (empty vector, as a control) (final concentration of 0.5 µg/µl) in OptiMEM and incubated for 30 minutes. The DNA:PEI complexes were added drop by drop to CC81LTRBLVGFP cells. Twenty-four hour post-transfection, fluorescence was observed using fluorescence microscopy and the percentage of GFP positive cells was determined by flow cytometry as described below.

### Transactivation of CC81LTRBLVGFP cells by valproic acid (VPA), trichostatin A (TSA) or INFα

CC81LTRBLVGFP cells were incubated for 24 hours to form a confluent monolayer. Subsequently, the cells were treated with 10mM VPA or 500nM TSA for 24 hours. To evaluate the effect of INF-α on BLV LTR promoter, CC81LTRBLVGFP cells were treated without or with 75, 125, 250, 500 or 1000 U/mL of INF-α for 48 hours. The percentage of GFP positive cells was determined by flow cytometry as described below.

### GFP protein expression analysis by flow cytometry

CC81LTRBLVGFP cells transiently transfected with pcDNABLVTAX-FLAG or pcDNA3.1 (empty) or treated with VPA or TSA were detached through trypsinization and transferred to a V-bottom plate. Subsequently, the cells were washed twice with PBS-BSA 3% and fixed with 4% PFA. Cell acquisition was performed using the Attune NxT (Thermo Fischer Scientific) cytometer, and the data was analyzed with the FlowJo software (Tree Star Inc.). Non-transfected or transfected with empty vector, and untreated cells were used as a control of the fluorescence background.

### Evaluation of the reporter cell line by infection with BLV

For setting up co-culture experiments, CC81LTRBLVGFP cells (as target cells) were co-incubated with FLKBLV producing cells (donors) at ratios of 1:1, 1:2, and 1:4 (donor:target) for 24 or 48 h at 37°C. Subsequently, the cells were washed with PBS, fixed with 4% PFA, and stained with DAPI (1/1000) for nuclei visualization. GFP expression was evaluated by means of an epifluorescence microscope and images were inspected using ImageJ. Once the experiment was set up, the co-culture was carried out by co-incubating 1,6x10^5^ CC81LTRBLVGFP cells with 0,4x10^5^ FLKBLV in a 12-well plate for 48 h at 37°C. The co-culture experiments using the BL3.1 cell line were performed as follows: CC81LTRBLVGFP cells were seeded at a density of 5x10^4^ cells per well in a 12-well plate and cultured at 37°C for 24 hours. Subsequently, they were co-cultured with BL3.1 cells at a density of 1x10^5^ cells per well for 72 h. For all experiments, co-culture cells were washed with PBS, fixed with 4% PFA, and stained with DAPI (1/1000) for nuclei visualization. GFP expression was evaluated by means of an epifluorescence microscope and images were inspected using ImageJ.

### Transwell infection assay

1,6x10^4^ FLKBLV cells, serving as viral particle-producing cells, were cultured on transwell inserts with a 0.4 μm pore size for 24 h to establish a monolayer. Following cell culture, the transwell inserts were incubated with CC81LTRBLVGFP reporter cells per well, which were previously plated in a 24-well. As a control, a similar number of FLKBLV and CC81 reporter cells were directly co-cultured in 24-well plates without an insert. The system was then incubated at 37°C for 48 h to facilitate the interaction between viral particles and CC81LTRBLVGFP cells. Cells were then fixed with 4% PFA, washed three times with PBS and subsequently blocked for 15min with PBS-3%BSA-0.1% Triton X-100. Cells were further incubated 1 h at RT with monoclonal antibodies anti-p24 (BLV3, 1/200, VMRD, USA). After three washes with PBS, cells were incubated for 1 h at RT with Alexa Fluor-594 conjugated goat anti-mouse IgG (Invitrogen) diluted 1/1,000 in PBS added with 3% (w/v) BSA and DAPI 1/1000, washed three times with PBS and mounted in 70% (v/v) glycerol pH 8.8. Cell infection was assessed using Zeiss LSM800 confocal microscopy.

### Fluorescence microscopy imaging

Mosaics of 5x5 tiles adding to twenty-five images per well were automatically acquired on Zeiss LSM800 confocal microscopy with a 10X objective. Two-source excitation was performed using 488 (power 0,92%) and 405 (0,86%) lasers, detection wavelength 510-700nm and 400-510nm respectively. Pinhole set at 25 μm. Images were taken at 1,03 μs pixel dwell time, a resolution of 4650x4650 pixel.

### Automated segmentation of fluorescently infected cells

A custom set of scripts was written in MATLAB to perform automated segmentation of nuclei and GFP+ cells and automated counting of nuclei per cell and classification into syncytia. The two emission channels (DAPI and GFP) were processed independently to segment objects based on the knowledge of the expected area of nuclei. In both cases, a pre-processing step was performed, involving brightness normalization to the [0.1-99.9]

% quantiles of the total brightness and gaussian filtering; for the GFP channel using a standard deviation of ¼ of the expected radius of nuclei and for the DAPI images a standard deviation of 0.1 of the expected radius of nuclei. Nuclei were segmented using a watershed-based algorithm with a size stopping rule at 4 times the expected area of a nuclei. GFP+ cells were segmented using simple Otsu thresholding allowing for a user-inputted factor to fine tune the segmentation. A post-filtering step was required to remove objects that were either smaller than twice the expected area of a nuclei or 200 time larger. Software is available upon request and will be uploaded to a public repository.

### Automated counting and feature measurement

Once segmentation step was completed in both channels, for each individual segmented GFP+ cell, the number of segmented nuclei within the same spatial region were counted, distinguishing between partially included and completely included accounting for segmentation errors. The nuclei number per GFP+ cell was established as the average between the total distinct nuclei inside and the total nuclei that were completely inside. The criteria to classify into syncytia was those GFP+ segmented cells with more than 3 detected nuclei. For completion a clustering step was performed using the distances between the centers of the detected nuclei, using a density-based algorithm (dbscan). The set of scripts allowed to automatically export a list of detected GFP+ cells (in the order of 300 per field of view), with their measured number of nuclei, relative brightness and measured area, and a list of detected nuclei (in the order of 10k per field of view). Software is available upon request and will be uploaded to a public repository.

## RESULTS

### Generation of a reporter cell line for BLV infection

A second-generation lentiviral system was utilized to develop the stable reporter cell line **Figure 1 A**. We replaced the CMV promoter in the pTripCMVGFP vector with the full-length 5’LTR of BLV to generate the reporter lentiviral vector. As shown in **Figure 1 B**, no GFP expression was detected for CC81 cells transduced with LVs LTRBLVGFP. This outcome is expected since the BLV LTR promoter has deficient basal activity and can only be activated and induce GFP expression in the presence of Tax or upon BLV infection. On the other hand, cell line expressing the BLV Tax transactivator protein (FLKBLV), transduced with LVs LTRBLV-GFP or CMV-GFP showed similar levels of GFP expression, indicating that the BLV promoter would have similar activity to the CMV promoter. Therefore, observing GFP in this cell line allows us to conclude that both the constructions and LVs production were successful. Finally, in both cell lines (CC81 or FLKBLV) transduced with CMVGFP LVs (control), GFP expression was observed. This result was anticipated, as these LVs containing the constitutive CMV promoter can induce high levels of GFP transgene expression.

**Figure 1.**
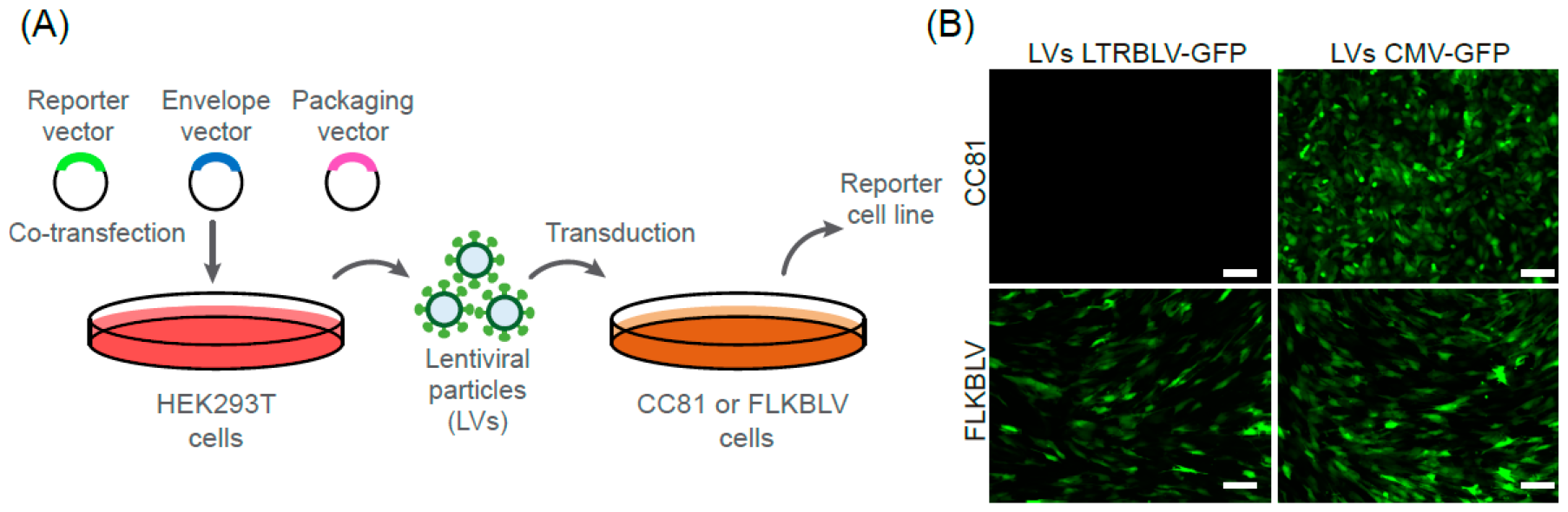
Generation of the reporter cell line for BLV infection. **(A)** Representative diagram of the process for obtaining the reporter cell lines CC81LTRBLVGFP and FLKBLVLTRGFP. HEK293T cells were co-transfected with packaging (DR8.74), envelope (VSV-G), and reporter (pTripLTRBLVGFP or pCMVGFP) plasmids. Forty-eight hours post-transfection, the supernatant containing lentiviral particles (LVs, LV-LTRBLVGFP or LV-CMVGFP) was collected. CC81 or FLKBLV cells were transduced with 250 μL of LV-LTRBLVGFP or LV-CMVGFP diluted in culture medium and 10 μL of polybrene 250 μg/mL and incubated for 60 h. **(B)** Representative images of transduced cell lines. The FLKBLV cell line served as a positive control. GFP expression was evaluated by fluorescent microscopy and the obtained images were analyzed using Image J. Scale bar, 100 μm.

### Functional testing of the reporter construct LTRGFP

To assess the effect of BLV Tax protein expression due to transactivation of the BLV LTR, CC81LTRBLVGFP reporter cells were transiently transfected with an expression vector pcDNABLVTAX-FLAG or the empty vector as a negative control. Tax expression in transfected cells was confirmed by immunofluorescence and western blot 24 h after transfection **(Supplementary Figure 1 A, B)**. We show that the induction of BLV Tax expression is enough to activate the BLV LTR in the reporter cell line, as evidenced by a significative increase of GFP+ cells. No GFP expression was detected in cells transfected with an empty vector or in untransfected cells **(Figure 2 A)**. The U5 region present in the LTR contains regulatory interferon factor binding sites, which have previously been reported to drive tax-independent replication of BLV [28]. We tested the effect of IFN-α treatment (0-1000 U/mL) on BLV LTR-driven GFP expression in the CC81LTRBLVGFP cell line. After 48 h of IFN treatment, we found no GFP expression in the cells at any of the concentrations tested **(Figure 2 B)**. Previous reports have shown that deacetylase inhibitors can induce Tax-independent transactivation of the LTR. For example, TSA or VPA has been shown to be an efficient activator of BLV expression [19, 26, 29]. We demonstrate that a small percentage of CC81LTRBLVGFP cells show GFP expression after treatment with TSA and VPA **(Figure 1 C)**. While there is a slight effect, it should be considered when designing experiments using our reporter cell line.

**Figure 2.**
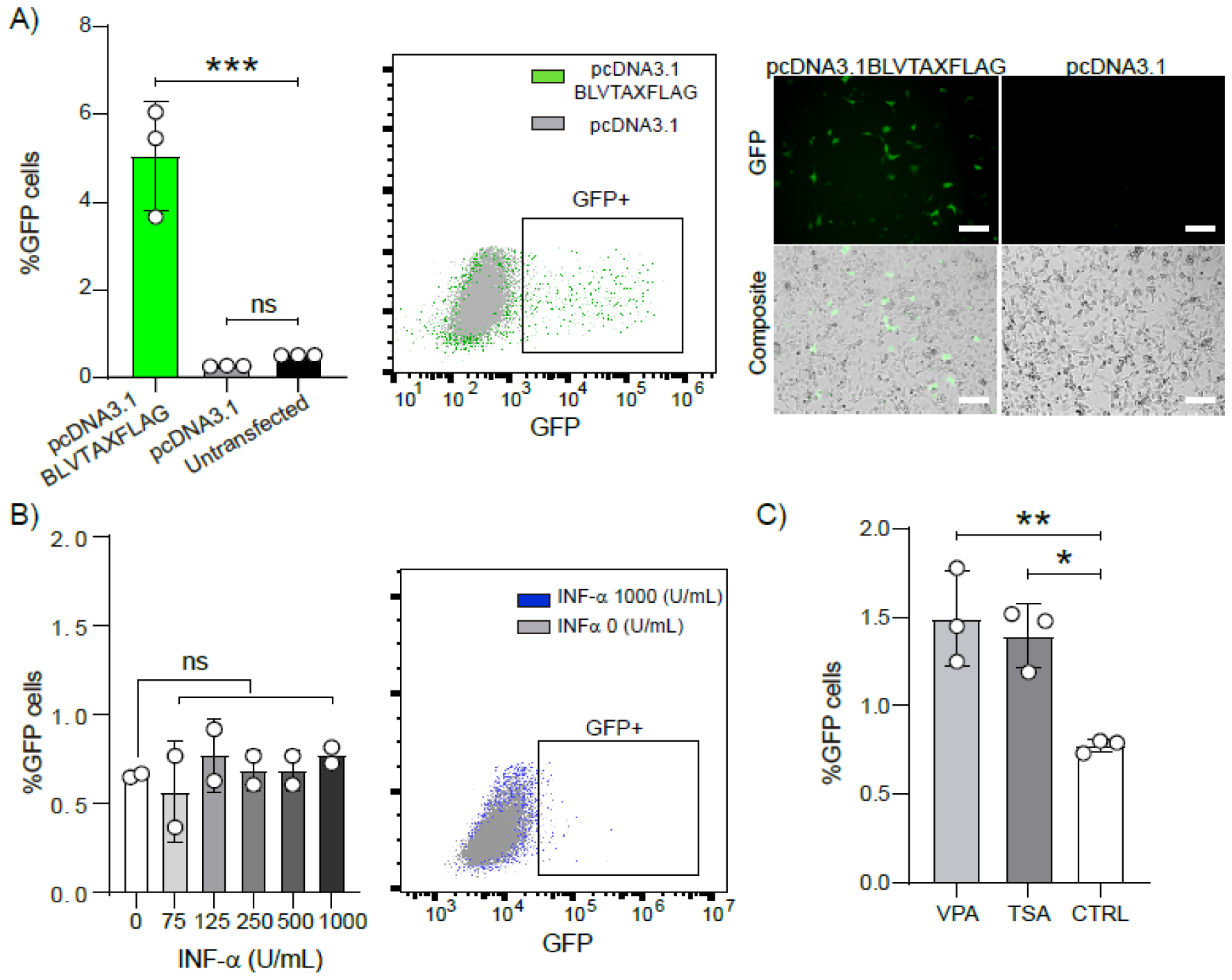
Functional testing of the reporter constructs LTRBLVGFP. **(A)** Transient transfection of the reporter cell line with Tax expressing vector. CC81LTRBLVGFP cells were transfected with the pcDNABLVTAX-FLAG or pcDNA3.1 empty vector (control). The %GFP+ cells were determined by flow cytometry. A representative dot plot of three replicates is shown. Bars represent the mean + SD of 3 replicates. ***p=0.0005, ns p=0.88 One-way ANOVA test. GFP expression was evaluated by fluorescent microscopy in CC81LTRGFP cells 48 h after transfection with Tax or the empty vector. Untransfected cells are shown as control. Scale bar, 100 μm. **(B)** The effect of INF-α on BLV LTR promoter activity was evaluated in CC81LTRBLVGFP reporter cells. Cells were cultivated for 48 h without or with 75, 125, 250, 500 or 1000 U/mL of INF-α, and the percentage of GFP+ cells was evaluated by flow cytometry. Bars represent the mean + SD of 2 replicates. One-way ANOVA test. **(C)** The effect of deacetylase inhibitors (VPA and TSA) on BLV LTR promoter activity was evaluated in CC81LTRBLVGFP reporter cells. Cells were cultivated for 24 h without (CTRL) or with 10 mM VPA or 500 nM TSA and the percentage of GFP+ cells were evaluated by flow cytometry. Bars represent the mean + SD of 3 replicates. **p=0.007 *p=0.01 One-way ANOVA test. For statistical analysis, GraphPad Prism 9.1.0 was used.

### Validation of the reporter cell line CC81LTRBLVGFP upon BLV infection

The next step was to evaluate the effect of BLV infection on LTR GFP expression by co-culturing the reporter cell line CC81LTRBLVGFP with persistently infected FLKBLV or BL3.1cell lines. We first optimized the co-culture between FLKBLV cells and the reporter cell line, assessing different ratios (1:1, 1:2, 1:4 FLKBLV:CC81LTRBLVGFP), and fluorescent syncytia (cell-cell fusion) formation was evaluated 24 and 48 h later by fluorescent microscopy. We determined that cell-cell fusion (the number of fluorescent syncytia) increases with the proportion of reporter cells to BLV-infected cells, particularly at the 4:1 ratio. Fluorescent syncytium can be detected 24 h after co-culture, but the number of syncytia and nuclei per syncytium increases at 48 h **(see Supplementary Figure 2)**. Thus, we conclude that the optimal conditions for co-culture between the FLKBLV and CC81LTRBLVGF cell lines involve a 1:4 ratio, respectively, maintained for 48 hours **(Figure 3, upper panel)**. Regarding the BL3.1 cell line co-culture, we first evaluated the same conditions used with FLKBLV (1:4 and 48 h). However, under these conditions, we did not observe the formation of fluorescent syncytia. Since BL3.1 cells grow in suspension, experimental conditions, such as the number of cells and the co-culture time were modified. Thus, we used 5x10^4^ CC81LTRBLVGFP cells per well (plated 24 h before co-culture), 1x10^5^ BL3.1 cells per well and 72 h co-cultured period. With this experimental set up, stronger fluorescence and larger syncytium were observed **(Figure 3, lower panel)**. In summary, our reporter cell line was able to express GFP and form syncytia when subjected to co-culture assays using cell lines persistently infected with BLV.

**Figure 3.**
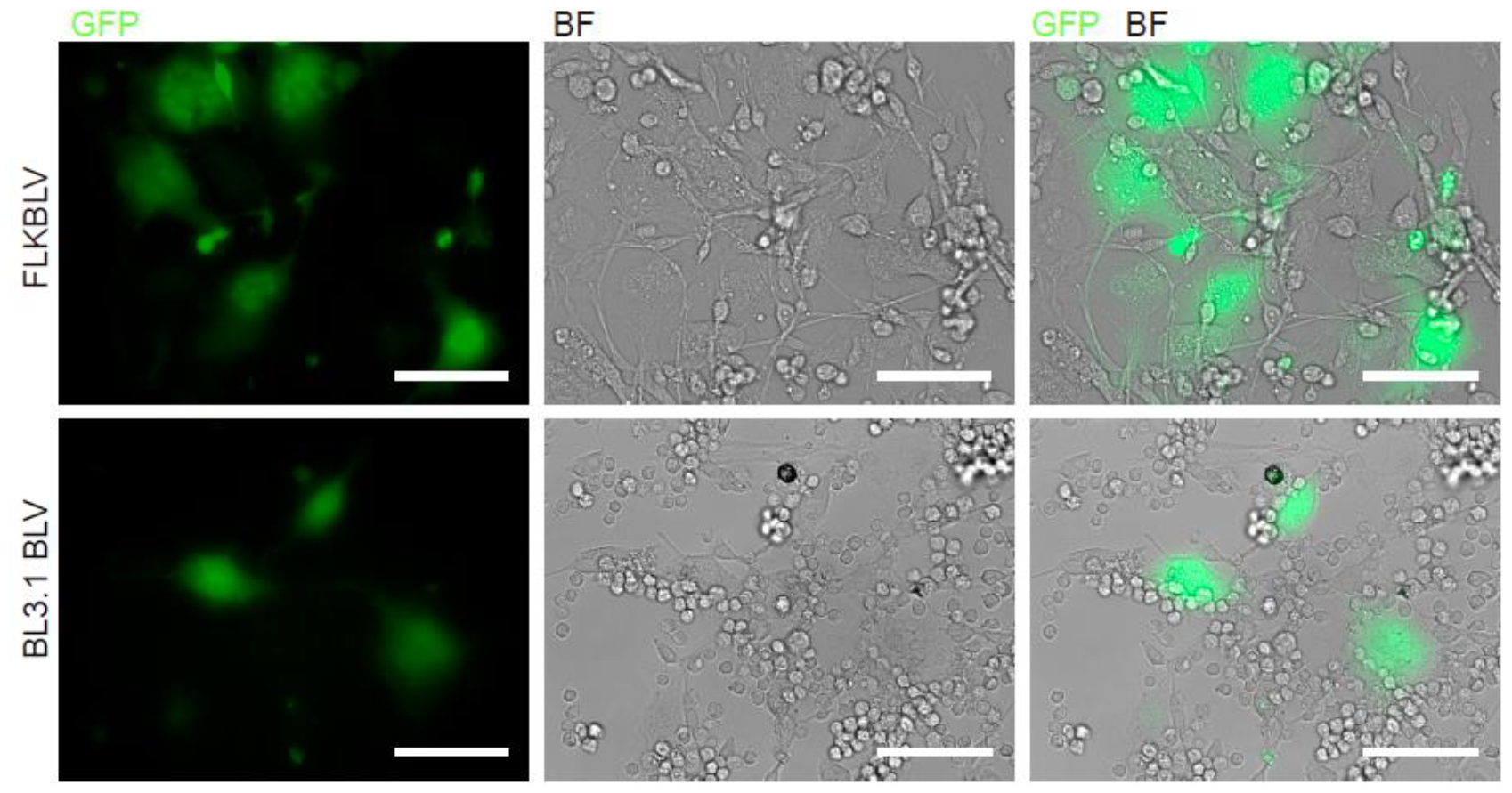
Co-culture of C81LTRBLVGFP reporter cell with the BLV infected cell lines FLKBLV o BL3.1. CC81LTRBLV cells were co-cultured in a 4:1 ratio with FLKBLV cells (above) or BL3.1 BLV cells (below) for 48 or 72 h, respectively. Syncytium expressed GFP was evaluated by fluorescent microscopy, and images were analyzed using Image J software. Representative images from three independent experiments are shown. BF: Bright field. Scale bar: 100 µm.

### CC81LTRBLV reporter cell system is useful to measure cell-to-cell (via cell-fusion) and cell-free BLV infection

To compare our BLV infection reporter system with traditional syncytia formation assay, CC81LTRBLVGFP reporter cells were co-cultured with FLKBLV cells in a 4:1 ratio for 48 h. BLV infection was measured concurrently by counting the number of syncytia using bright-field microscopy and the number of GFP+ syncytia using fluorescent microscopy in the same field. We found a significative correlation between the GFP syncytia number using the reporter cell line and the syncytia number assessed by the traditional syncytia formation assay (Pearson correlation coefficient, r= p<0.0001) **(Figure 4 A, Supplementary Table 1)**. However, with our reporter cell line, we were able to distinguish GFP+ syncytia that were not detected with the traditional method (˃30%). In addition, since traditional syncytia formation assays consider multinucleated cell with 4 or more nuclei, we detected individual infected cells and multinucleated cells with fewer than 4 nuclei (herein termed “GFP+ infected units”). Taking these results into consideration, we utilized an automated image acquisition system (25 images per well) using confocal microscopy and developed a software that allows for detection and counting of GFP+ infected units. This automated method allows detection of individual infected and multinucleated cells with less than 4 nuclei (30% of total counts) indetectable by traditional method **(Figure 4 B, Supplementary Figure 3, Supplementary Table 2)**. To test the ability of CC81LTRBLVGFP cells to detect BLV cell-free infection, we initially used FLKBLV supernatant containing BLV viral particles to infect the cell line; however, no GFP+ cells were observed. To further investigate the ability of our reporter cell line to detect cell-free BLV infection, we performed a transwell experiment between the CC81LTRBLVGFP reporter cells and an insert with FLKBLV cells producing BLV particles and immunofluorescence. Here, we detected CC81 reporter cells infected with BLV by detecting the capsid protein and individual cells with a strong GFP signal using fluorescent microscopy **(Figure 4C)**. The number of GFP+ cells is lower than when cell contact is allowed.

**Figure 4.**
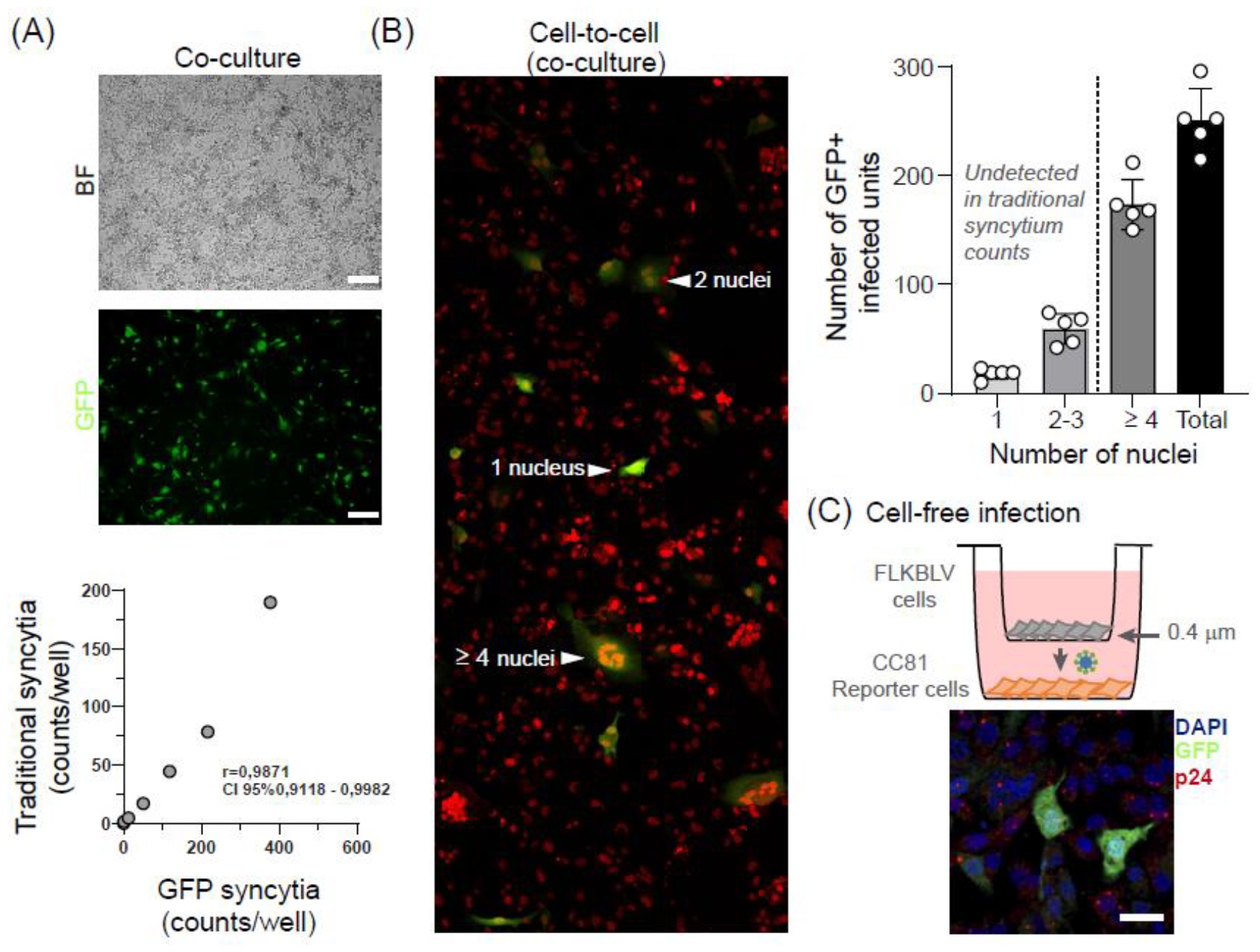
CC81LTRBLV reporter cell system is useful to measure cell-to-cell (via cell-fusion) and cell-free BLV infection. **(A)** The correlation between the stained syncytia counts in traditional assay and the fluorescence syncytia counts. 5x10^4^ CC81LTRBLVGFP cells were co-cultured with 0, 90, 180, 750, 1500, 3000 and 6000 FLKBLV cells in a 24-well plate at 37ºC for 48 h after that, the cells were fixed with 4% PFA. The number of syncytia per well, defined as multinucleated cells with 4 or more nuclei, was counted (in the same field) by two independent experimenters in bright field (BF) and fluorescence images. Syncytia count measured by the traditional syncytia assay was plotted against syncytia number determined by GFP expression and the correlation was evaluated with the Pearson correlation test, r=0.9871. Scale bar 200 μm. **(B)** A representative section of an image acquired automatically obtained from the co-culture between CC81LTRBLVGFP cells and FLKBLV cells is shown. GFP+ cells with 1, 2 or more than 4 nuclei are indicated (white triangles). DAPI staining is shown in red. The number of GFP+ cells per well was measured with 1, 2-3, 4 nuclei, and total. Bars represent the mean + 3SD of five independent experiments. **(C)** A BLV cell-free infection experiment was performed using the transwell system. After 48 hours of culture, the cells were fixed with 4%PFA and subjected to immunofluorescence with monoclonal anti-p24 (to detect BLV capsid protein, in red). Cell nuclei were stained with DAPI (blue). Cell infection was assessed using Zeiss LSM800 confocal microscopy. Scale bar, 25 μm.

## DISCUSSION

It is well established that purified BLV virions exhibit inefficient infection of susceptible cells and transmission in cell culture is greatly improved when cell-to-cell contact is allowed. When BLV-infected cells are co-cultured with susceptible cells, they undergo cell-to-cell fusion events (syncytia) in cell culture [18], [16]. This syncytium formation assay is a primary tool in studying BLV infectivity [15, 17, 30]. Although this method is highly robust, the complexity of visually counting syncytia poses a significant technical challenge, especially in high-throughput screening assays.

An alternative to this challenge is the development of reporter cell lines capable of emitting a fluorescence or luminescence signal in response to viral infection. In this sense, an improved reporter cell system utilizing the BLV full-length LTR [22] or LTR/U3 promoter [19–21] was described. Recently, Sato *et al*., [21] established a reporter cell line, namely CC81-GREM, harboring the EGFP reporter gene under the control of the GRE-mutated LTR-U3 promoter. This assay was used to evaluate the BLV-infected white blood cells, the capacity of neutralizing antibodies, and the infectious potential of cells in milk from BLV-infected dams, demonstrating greater sensitivity in quantifying BLV infectivity than traditional cell lines used in conventional cell fusion assays.

In this study, we constructed a reporter cell line to measure BLV infectivity using a lentiviral vector system due to the ease of establishing stable cell lines [31]. This cell line has stably integrated into its genome a construct containing the viral BLV full-length 5’LTR associated with the fluorescent protein GFP coding region. The LTR promoter is activated in the presence of the viral protein Tax (transcriptional activator), which is only present when viral infection of the reporter line occurs.

We first characterize the functionality of our reporter cell line through Tax-dependent and Tax-independent transactivation. As previously described, the 5’LTR region present in the BLV genome contains protein binding regulatory sites and is responsible for virus integration and replication. This mechanism is significant because the BLV LTR-U3 regulates BLV replication induced by Tax protein [32, 33]. In this regard, through transient transfection of the CC81LTRBLVGFP reporter cell line with a Tax expression plasmid, we demonstrated that the Tax protein efficiently activated the CC81LTBLVGPF cell line, observing cells expressing GFP. This result is consistent with previous studies [22], [20]. On the other hand, it has been demonstrated that using HDACi can induce the expression of both viral and cellular genes. In this context, previous studies using reporter-based assays have shown that BLV expression is upregulated in response to TSA and VPA stimulation in a Tax-independent manner [26]. In this sense, as expected, low levels of GFP expression were observed when stimulating our reporter cell line with VPA and TSA.A binding site for interferon (IFN) regulatory factors 1 and 2 (IRF-1 and IRF-2) has been identified in a transcriptional enhancer in the LTR-U5 region, and the possibility of BLV LTR basal activation by IFN in a Tax-independent manner has been suggested [28]. One of the questions we asked ourselves was if stimulation of the reporter cell line with IFN was sufficient to activate the promoter. We showed that high concentrations of IFN do not induce GFP expression, suggesting that our reporter cell line (containing the complete LTR) could be used to study the effect of IFN at different stages of BLV infection.

We validated the CC81LTRBLVGFP reporter cell line following BLV infection by assessing its ability to form syncytia through cocultivation with FLKBLV or BL3.1 cell lines persistently infected with BLV. The quantification of fluorescent syncytia using CC81LTRBLVGFP strongly correlated with traditional syncytium counts (r=0.9871). However, when the reporter cell line was cocultured with FLKBLV, we observed both single GFP+ cells and fluorescent multinucleated cells with 2 or 3 nuclei. These observations suggest that CC81LTRBLVGFP can detect infection events that are not distinguishable in traditional syncytium formation assays improving the robustness of our method.

In this regard, we integrate automated scanning and image acquisition using a confocal microscopy, with the development of an analysis software that allows us to detect and count single GFP-positive cells or fluorescent multinucleated cells with 2 or more nuclei. This approach significantly accelerates sample analysis compared to traditional visual counting methods. Through coculture assays between CC81LTRBLVGFP and FLKBLV, we observed that BLV predominantly spreads through cocultured cells via cell fusion transmission. Additionally, we evaluated the usefulness of the reporter cell line in measuring cell-free infection through a transwell assay. We successfully detected GFP cells and viral particles within the reporter cell line by labeling a BLV capsid. However, we cannot exclude the possibility that Tax is secreted and can enter CC81 LTRBLVGFP to transactivate the LTRGFP. Furthermore, Tax has been identified in exosomes released from cells infected with HTLV [34]

It has been previously documented that retrovirus spread in cultured cells and tissues via two routes: through cell-to-cell contact and cell-free mode through the extracellular environment and infecting new cells [13, 35–37]. Cell-to-cell transmission typically involves tight cell-cell contact, such as virological synapses, biofilms, or the formation of nanotubes. On the other hand, cell-cell fusion has been identified as an alternative pathway for cell-to-cell transmission [12, 38]. Cell-cell fusion has been described in HIV and HTLV. However, the relevance of this property for virus spread *in vivo* is unclear, and there is no clear evidence that syncytia formation can enhance HIV-1 dissemination *in vivo* [13].BLV is naturally and mainly transmitted via cell-to-cell rather than cell-free mechanisms, as the extent of cell-free infectivity of virions in the blood of BLV-infected cattle is significantly lower [39]. Currently, very little is known about the mode of transmission between cells or the efficiency of cell-free infection for BLV. Our results support that cell-to-cell transmission is the primary mechanism for BLV spread in cell co-culture. In addition, we observed that more than 90% of infection events (GFP+ infected units with ˃1 nuclei) involved cell-cell fusion. In this regard, our reporter cell line, combined with the development of analysis software, is a useful tool for understanding the role of cell fusion in BLV transmission.

Although the co-culture of infected and target cells is a frequently used and efficient method for studying BLV infection, some concerns remain. For example, it is challenging to distinguish between donor and acceptor cells. Cell-free BLV infection models allow for more controlled experimental conditions. Further exploration of the mechanisms involved in cell-to-cell or cell-free transmission of BLV is necessary to better understand its pathogenesis and biology. Systems that can measure both mechanisms of infection, like our reporter cell line, could be handy for this.

Finally, our reporting system could be implemented to evaluate the infective potential of BLV present in different biological samples of importance in the veterinary management of cattle, such as colostrum, semen, mucous fluids, and more. Additionally, it could be used to assess the effect of different disinfectant measures on infectivity, such as heat treatment, specific disinfectants, and other methods of viral inactivation. Lastly, it could also be employed in the evaluation of molecules with antiviral activity (e.g., BLV protease inhibitor compounds) or molecules capable of neutralizing infection (e.g., specific monoclonal antibodies and their smaller derivatives).

## AUTHOR CONTRIBUTIONS

NO-D and OP: conceptualization and designed the project; NO-D and FR designed the experiments; NO-D and FR performed the experiments; NO-D, FR, MF, FC, CO and FDN analyzed the data, AV: design image processing software. All authors contributed to the preparation of the manuscript.

## ACKNOWLEDGMENTS

Special thanks to Mabel Berois for insightful discussion and critical reading the manuscript. We thank Marcela Díaz from the Advanced Bioimaging Unit at the Institut Pasteur Montevideo & Universidad de la República, for her technical assistance in acquiring confocal microscopy images. The authors gratefully acknowledge the Cell Biology Unit at the Institut Pasteur Montevideo for their support & assistance in the present work.

## ETHICAL APPROVAL

This article does not contain any studies with animals performed by any authors.

