## Supplemental Figures for "A versatile reporter system to study cell-to-cell and cell-free bovine leukemia virus infection"

### *Supplementary Material*

#### *Supplementary Materials and Methods*

**Western blotting.** In brief, transfected CC81LTBLVGFP HEK293T cells were collected and lysed in RIPA buffer (50 mM Tris, pH 7.5, 150 mM NaCl, 1 mM 100 EDTA, 1% Nonidet P-40, 0.1% SDS, protease inhibitor cocktail) for 30 min on ice, followed by centrifugation for 10 min, 12,000 x g at 4°C. Cell lysate then boiled at 100 °C for 10 min with 1XSDS loading buffer containing 2-Mercaptoethanol. Samples were run on 10% SDS-PAGE gels, transferred to PVDF membranes, FLAG-tagged Tax was detected using a primary mouse anti-FLAG (1/3000) (Sigma#F1804), followed by anti-mouse HRP (1/5000). Anti-actin (1/3000, Sigma#A3853) was employed as a control. Membranes were incubated with the enhanced chemiluminescence enzyme substrate kit (Thermo Fisher Scientific, USA), analyzed by Image Quant (GE).

**Immunofluorescence Assay.** The expression of Tax proteins was evidenced by indirect immunofluorescence and confocal microscopy. Briefly, 50 µl of transfected cells were fixed for 30min with 4% paraformaldehyde, washed three times with PBS, and subsequently blocked for 15min with PBS-3%BSA-0.1% Triton X-100. Cells were further incubated 1 h at RT with primary mouse anti-FLAG (1/500) (Sigma#F1804). After three washes with PBS, cells were incubated for 1 h at RT with anti-mouse Alexa Fluor 594 (1/1000) (Invitrogen) in PBS added with 3% (w/v) BSA and DAPI 1/1000, washed three times with PBS and mounted in 70% (v/v) glycerol pH 8.8. Images were generated using an epifluorescence microscope (Olympus IX81) and were analyzed using ImageJ software.

*Supplementary Figures*

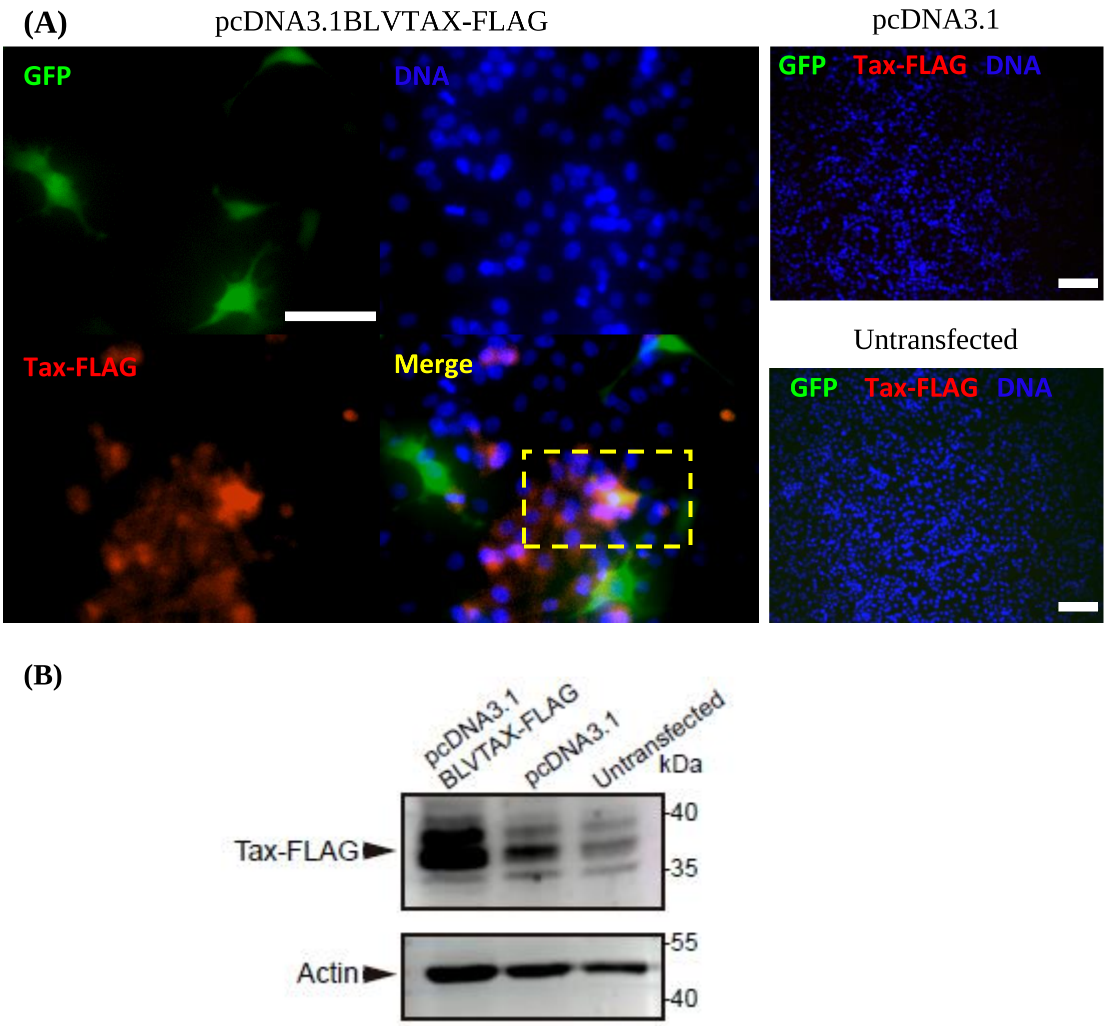


**Supplementary figure 1. Expression of BLV Tax protein in CC81LTRBLVGFP reporter cell line.** CC81LTBLVGFP were transitory transfected with expression plasmid pcDNA3.1BLVTAX-FLAG or pcDNA3.1 (empty vector) (2.5 ug) and Tax expression was evaluated by indirect immunofluorescence or western blot. **(A)** After 24 h, cells were fixed and permeabilized with 4% PFA and 0.1% Triton X, respectively, and stained with primary mouse anti-FLAG (1/500) (Sigma#F1804), followed by anti-mouse Alexa Fluor 594 (1/1000). Staining of nuclei was performed using DAPI (1/1000). Images were generated using an epifluorescence microscope (Olympus IX81). GFP (green), Tax-FLAG (red), DNA (blue) and merge (yellow). Scale bar, 50 µm (pcDNA3.1BLVTAX-FLAG). Scale bar, 100 µm (pcDNA3.1 and untransfected). **(B)** After 24 h, cells were lysed, and extract obtained was analyzed by Western blot. FLAG-tagged Tax was detected using a primary mouse anti-FLAG (1/3000) (Sigma#F1804), followed by anti-mouse HRP (1/5000, Invitrogen). Anti-actin (1/3000, Sigma#A3853) was employed as a control. Protein sizes were indicated by Page Ruler Prestained Protein Ladder, 10–180 kDa (Thermo Fisher Scientific). Untransfected cells were used as a control.

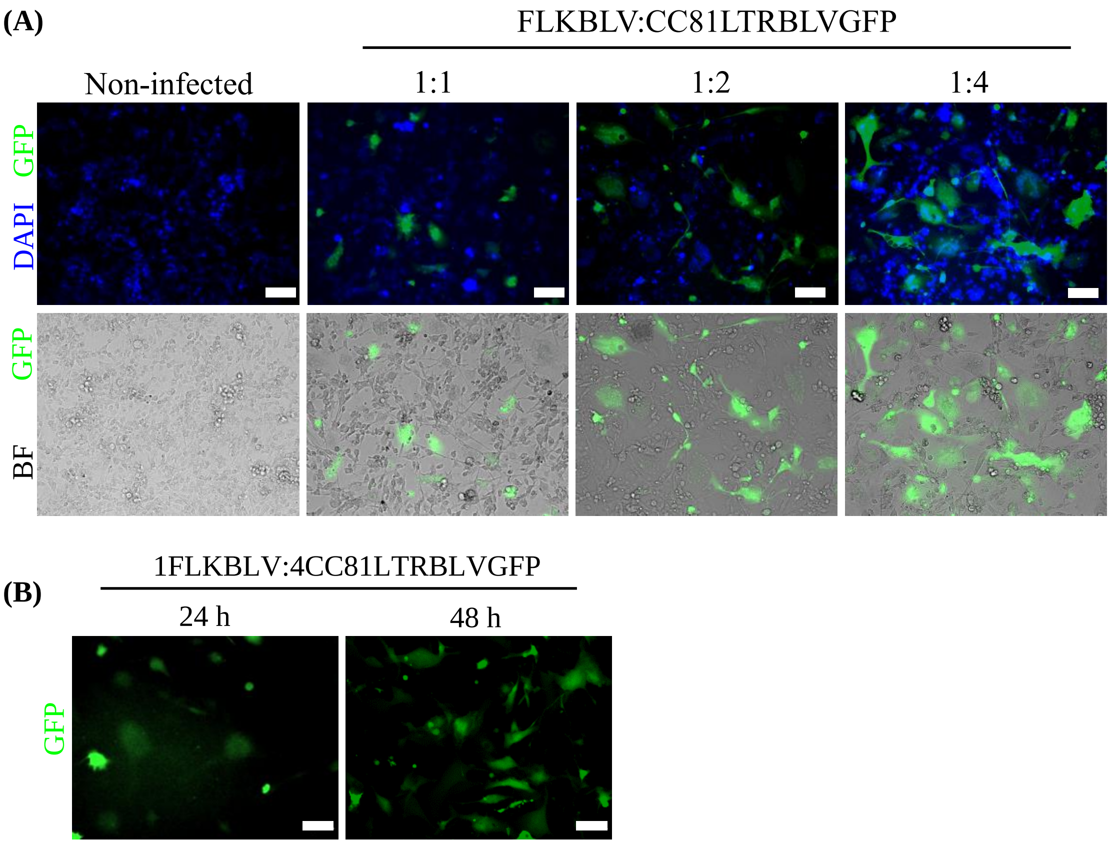


**Supplementary figure 2. Co-culture assay optimization. (A)** Co-culture was performed between the FLKBLV cell line and CC18LTRBLVGFP reporter in the ratios 1:1, 1:2 and 1:4. At 48 h post co-culture, cells were fixed with 4 % PFA and nuclei stained with DAPI (1/1000). Images were taken under the microscope of epifluorescence. The images were captured using an epifluorescence microscope. Scale bar, 100 μm. **(B)** Co-culture was performed between the FLKBLV cell line and CC18LTRBLVGFP reporter in the ratios 1:4 and the appearance of fluorescence was evaluated at 24 and 48 h**.**

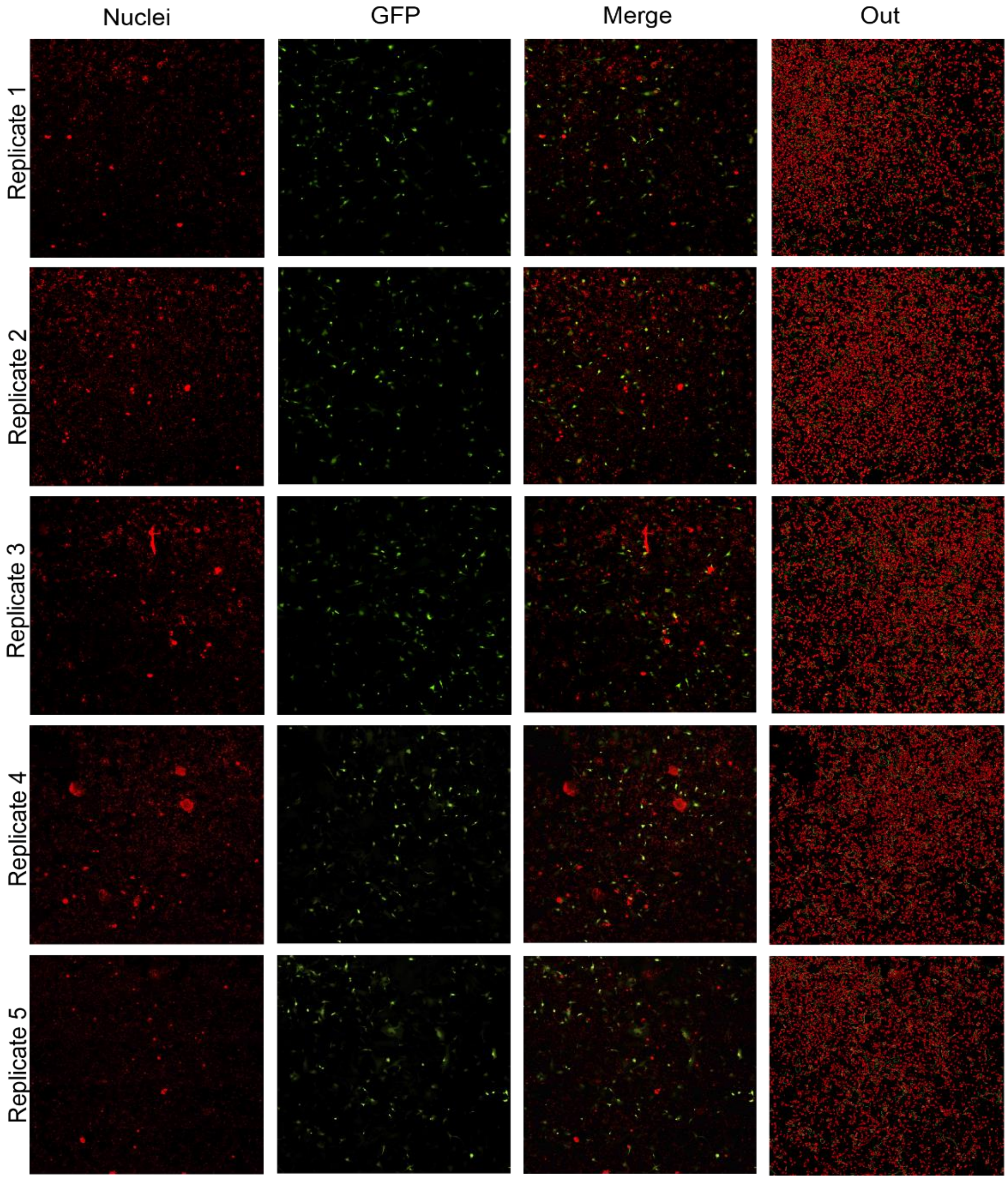


**Supplementary figure 3. Five replicates of automated counting analysis of fluorescently infected cells.** Mosaics of 5x5 tiles adding to twenty-five images per well were automatically acquired on Zeiss LSM800 confocal microscopy with a 10X objective. Automated segmentation of fluorescently infected cells was carried out and Automated counting and feature measurement.
